# Multimodal neuroimaging markers of variation in cognitive ability in older HIV+ men

**DOI:** 10.1101/2020.11.26.399592

**Authors:** Ana Lucia Fernandez Cruz, Chien-Ming Chen, Ryan Sanford, D. Louis Collins, Marie-Josée Brouillette, Nancy E. Mayo, Lesley K. Fellows

## Abstract

**Objective:** This study used converging methods to define the structural and functional characteristics of the neural substrates underlying variation in cognitive ability in older men with well-controlled HIV infection.

**Methods:** Seventy-six HIV+ men treated with combination antiretrovirals completed attention and inhibitory control tasks tapping different cortico-subcortical circuits while time-locked high-density EEG was acquired. Fifty-four also underwent structural MRI. We investigated relationships between task-evoked EEG responses, cognitive ability and immunocompromise. MRI suggested a subcortical basis for the observed EEG effects.

**Results:** EEG activity was associated with cognitive ability at later (P300) but not earlier processing stages of both tasks. However, only the P300 evoked by the attention task was associated with past HIV infection severity. Source localization confirmed that the tasks engaged different brain circuits. Thalamus volumes correlated with P300 amplitudes evoked by the attention task, while globus pallidus volumes were related to the P300 in both tasks.

**Interpretation:** This is the first study to combine structural and functional imaging in an overlapping sample to address the neural circuits related to cognitive dysfunction in HIV. Neural substrates of attention were more affected than those supporting inhibitory control. Preliminary evidence suggests these differences may relate to vulnerability of the thalamus to the effects of HIV. Our results suggest high-yield tasks and circuit targets for future work.

## Introduction

Despite effective viral suppression, cognitive impairment remains common in persons living with HIV, particularly as they grow older, with prevalence as high as 40% (Heaton et al., 2010; Su et al., 2015). Executive function, attention, and processing speed are commonly affected (Goodkin et al., 2017; Valcour et al., 2004). While typically mild in people taking combined antiretroviral therapy (cART), impairments can nonetheless limit everyday function (Becker, Thames, Woo, Castellon, & Hinkin, 2011; Heaton et al., 2004; Mayo et al., 2019a). The causes and underlying neural mechanisms remain unclear, with no consensus on optimal neuroimaging markers or intervention targets.

MRI studies have reported regional brain volume loss affecting subcortical structures and cortical regions in people with HIV compared to HIV-controls (Holt, Kraft-Terry, & Chang, 2012; Sanford et al., 2016; Thompson & Jahanshad, 2015). However, the patterns of structural differences and their relationships with cognitive impairment have been inconsistent across studies (Chang & Shukla, 2018; Masters & Ances, 2014). Whether brain injury in HIV and the resulting cognitive impairment are the manifestations of a generalized process affecting the whole brain or due to specific dysfunction in vulnerable regions or circuits has yet to be established.

Structural imaging alone is unlikely to provide the answer. In contrast to neurodegenerative disorders such as Alzheimer’s disease, many of the candidate pathophysiological mechanisms in HIV may not cause frank neuronal loss. Local cytokine release due to neuroinflammation, for example, might impair synaptic plasticity or interfere with neurotransmission, without yielding macroscopic brain atrophy (Nolan & Gaskill, 2019). EEG (and magnetoencephalography (MEG)) offer insights into brain function. Paired with appropriate cognitive tasks, these methods can probe specific circuit function with excellent temporal resolution. EEG is non-invasive and relatively inexpensive, so could be made widely available, even in resource-poor settings, yet there has been surprisingly little application of these methods in HIV (Fernández-Cruz & Fellows, 2017). There is evidence, albeit mainly from small sample studies, that EEG/MEG measures can distinguish people with HIV infection from HIV-controls, and HIV+ individuals with and without HIV-associated neurocognitive disorders (HAND) (Fernández-Cruz & Fellows, 2017; Groff et al., 2020; Lew et al., 2018; Wiesman et al., 2018).

Here, we recorded stimulus-locked high-density EEG during performance of two demanding test of executive function: an inhibitory control task (Simon task) and an attention and working memory-requiring categorization task (auditory oddball) in the same sample of older men with well-controlled HIV infection, a sub-sample of whom also underwent structural MRI. In both tasks, we distinguished earlier EEG activity reflecting initial cortical sensory processing and conflict detection and later activity reflecting higher-order cognitive processes (Luck, Woodman, & Vogel, 2000).

We aimed to provide evidence of the neural mechanisms underlying variation in the mild cognitive impairment to normal range, addressing specificity at the level of neural circuits and stages of cognitive processing, i.e., earlier vs. later brain activity. Structural MRI was used to localize cortical sources of EEG activity and to explore whether EEG results were explained by structural variation in subcortical nuclei implicated in these EEG responses.

## Materials and Methods

### Participants

Eighty-nine people living with HIV drawn from the Positive Brain Health Now (BHN) cohort participated in this study. BHN is a longitudinal study of brain health in people over age 35 living with HIV in Canada, on cART. Exclusion criteria were dementia severe enough to preclude informed consent (Memorial Sloan Kettering severity ≥ 3 (Price & Brew, 1988)), current substance dependence or abuse, psychotic disorder or non-HIV related neurological disorder, or terminal illness. Demographic and clinical data were acquired by self-report questionnaires, chart review and a brief cognitive test battery (M. J. Brouillette, Fellows, Finch, Thomas, & Mayo, 2019; Mayo, Brouillette, & Fellows, 2016). We acquired high-density EEG in all participants as part of the baseline assessment prior to randomization into two intervention sub-studies (trials of cognitive training or physical exercise) that sampled from the BHN cohort (**Fig. 1**). These sub-samples were randomly drawn from BHN participants in Montreal, based on scores on the cognitive assessment battery. 69 participants were drawn from those scoring below the median, 20 from those above the median. Structural MRI was also available in the 58 participants enrolled in the cognitive training study (Mayo et al., 2016). The protocol was approved by the ethics board of the McGill University Health Centre and all participants provided written informed consent.

**Figure 1:**
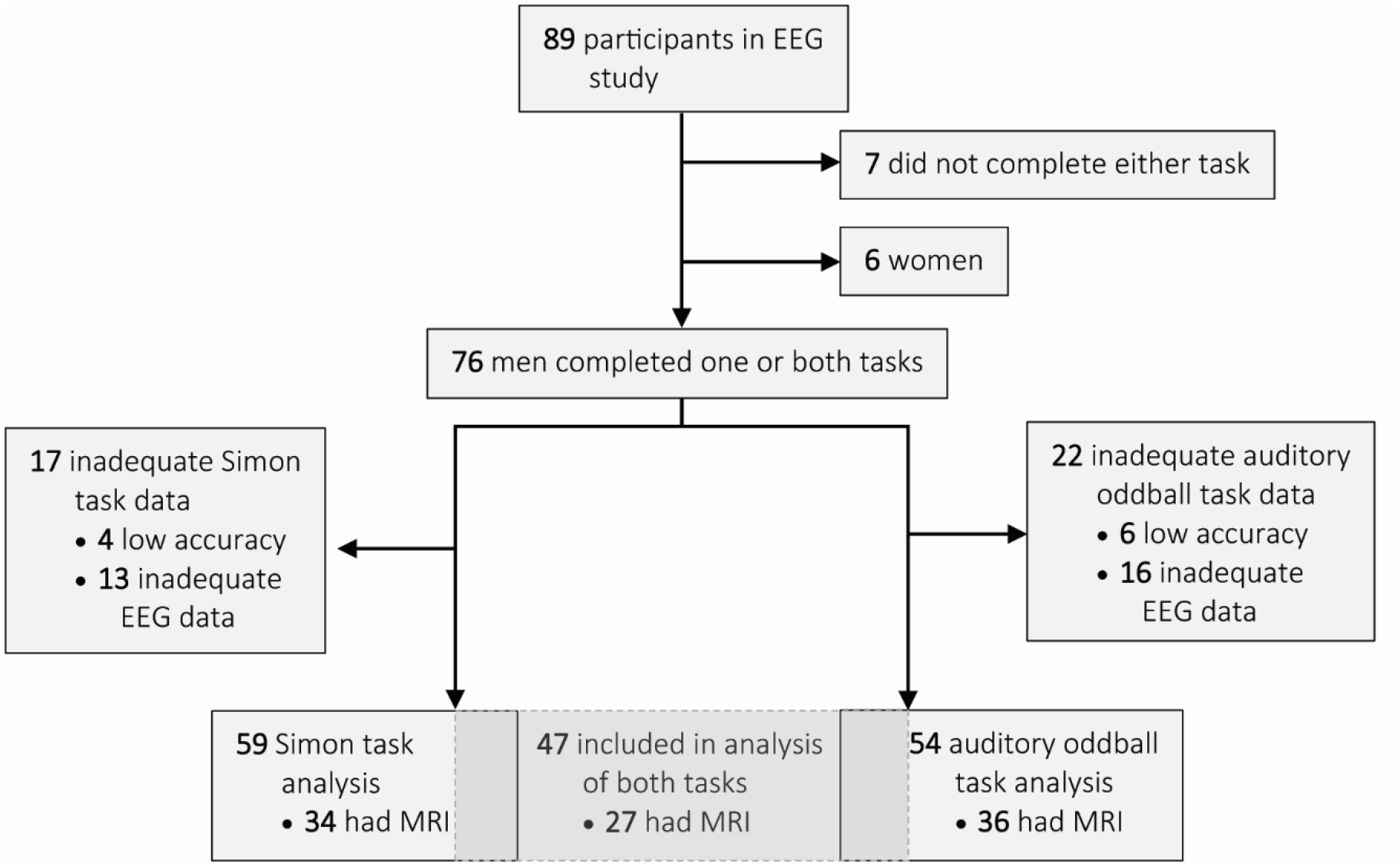
CONSORT diagram of the study. Participants who performed the task at chance accuracy or who had < 25 usable EEG trials after pre-processing were excluded. Data were only available for 6 women, insufficient to reliably adjust for sex effects, so the analysis was restricted to men. Sixty-six men had complete data for at least one task, 47/66 had complete data for both tasks. Structural MRI was acquired in 43/66. Brief cognitive assessment was done in all participants.

### Cognitive and Electrophysiological Assessments

#### Auditory Oddball Task

The multiple auditory oddball task is shown in **Figure 2**. Audiometry was carried out to ensure hearing adequate to perform the task. On each of 260 trials, participants fixated a cross on the computer screen and heard a series of 4 tones, the third of which might differ in frequency slightly (small deviant) or substantially (large deviant) from the other three. Participants reported whether this tone was the same or different with a left- or right-hand button press. Participants had a practice block with 28 trials before EEG recording began. The sensitivity index *d’* was calculated for the large and small deviant tones with the formula *d′* = *hit rate* − *false alarm rate* to estimate discrimination accuracy.

**Figure 2:**
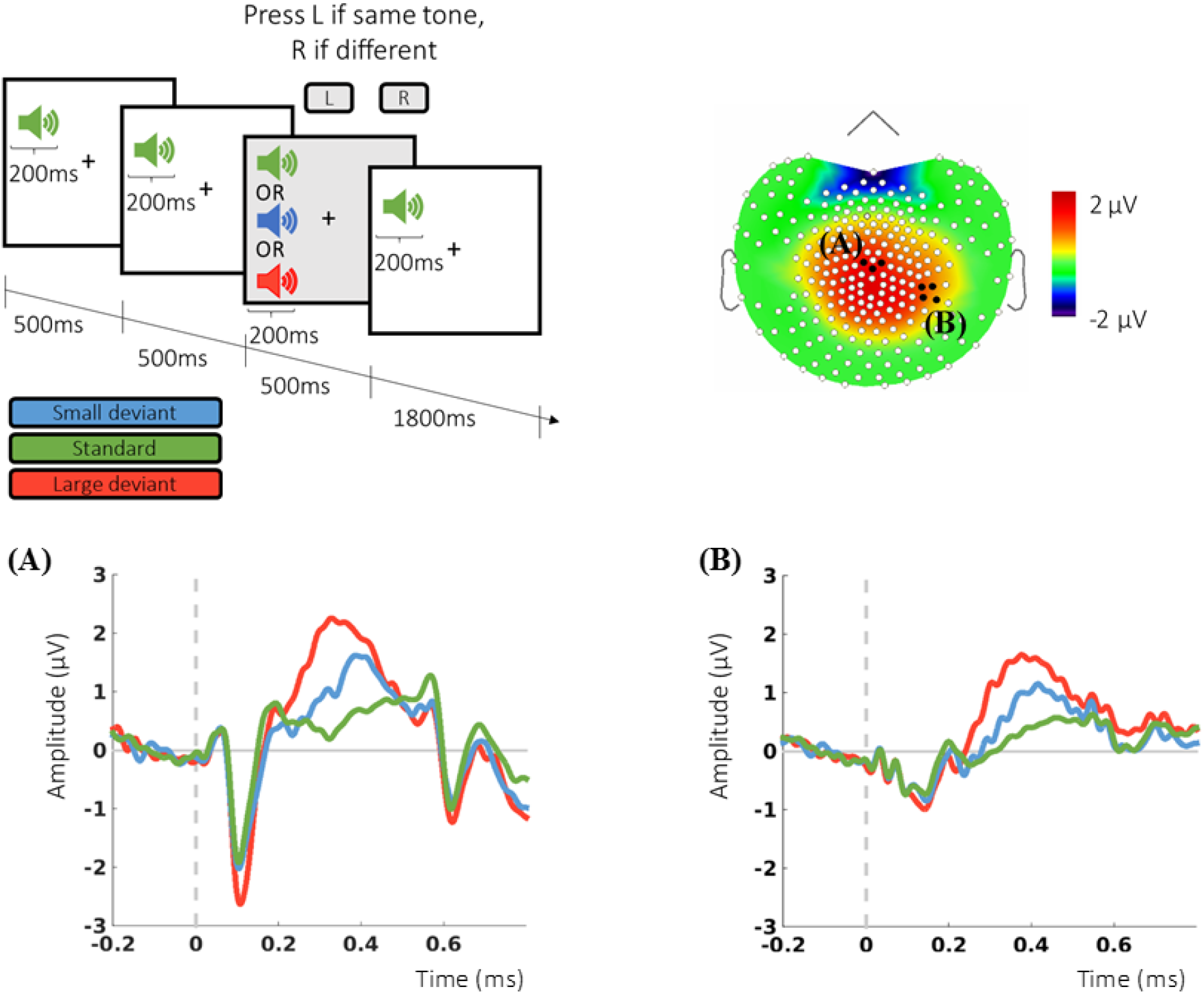
Oddball task and evoked potentials. Upper panel shows an example trial of the multiple auditory oddball task and the grand average topographic maps for the P300 with the 2 clusters of electrodes used for analysis highlighted (A and B). Participants completed 260 trials, each consisting of a sequence of four 200-milliseconds long tones presented binaurally at 70dB. The first, second and last tone of each of the trials were always “standard” tones (1047 Hz), the third tone could be either a standard tone or one of two deviant tones; a “small deviant” tone (1078 Hz) or a “large deviant” tone (1175 Hz). Participants were instructed to fixate a cross in the middle of the computer screen, and to report whether the third tone of each of the series was the same (right button press) or different (left button press) from the other three tones. Of the 260 trials, 146 were standard trials, 57 were small-deviant trials and 57 were large-deviant trials, presented in random order. **(A)** Grand average over the electrode cluster surrounding Cz and **(B)**over a cluster chosen following a data-driven approach.

#### Simon Task

The Simon task required participants to make a left or right button press response according to an arbitrary sensorimotor rule: one of two shape cues was associated with the left button, the other with the right button (randomized across participants). The shape appeared on the left or right side of the screen, inducing a pre-potent response tendency to respond with that hand. The cue and side of presentation matched on congruent trials (190) and conflicted on incongruent trials (190) (**Figure 3).** Sixteen practice trials preceded the main task. Reaction time (RT) and accuracy were recorded.

**Figure 3:**
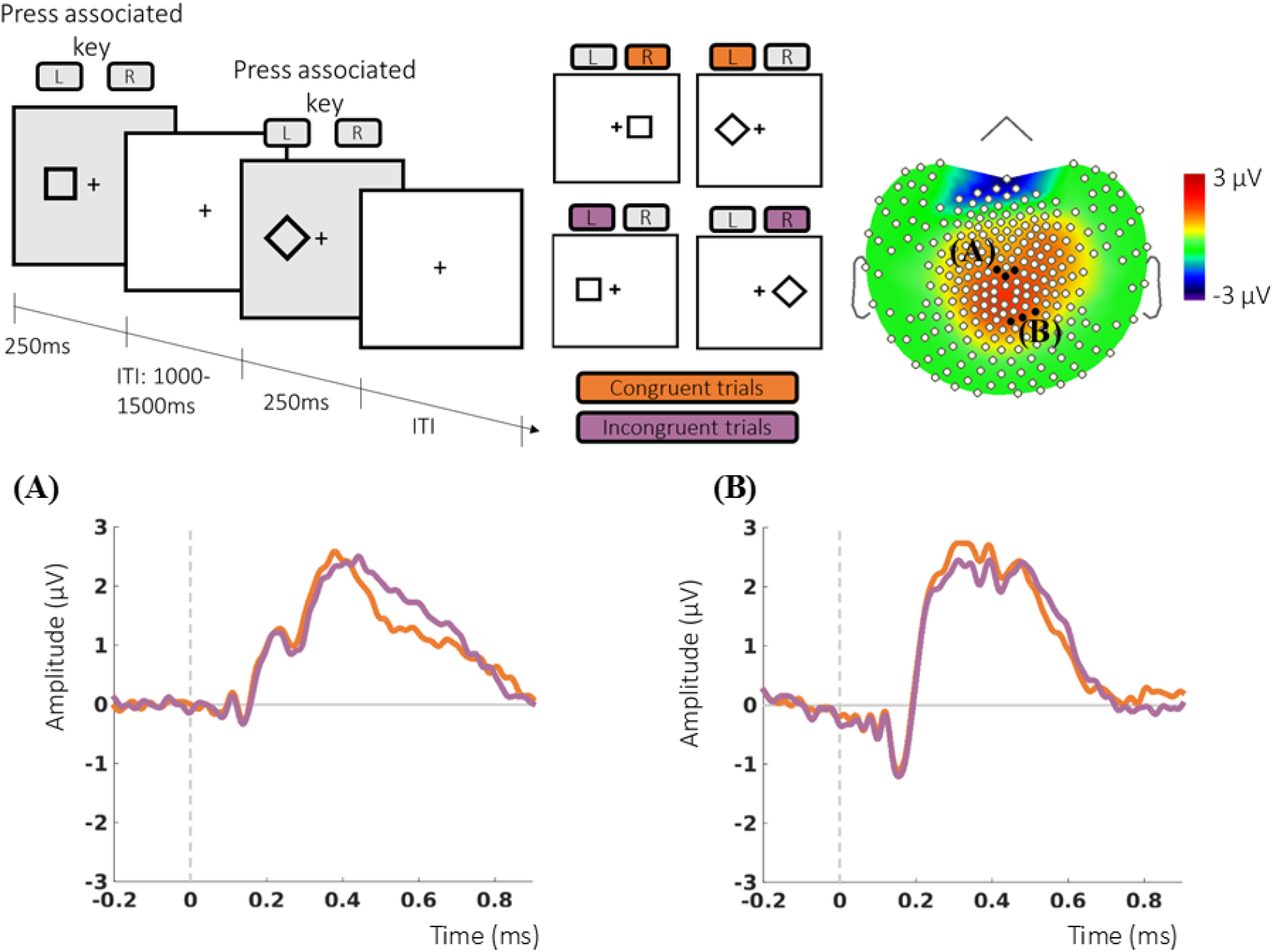
Simon Task and evoked potentials. Upper panel shows example trials of the Simon task and the grand average topographic maps for the P300 with the two clusters of electrodes used for analysis highlighted (A and B). During the Simon task (upper right panel) participants responded to one of two 2.5 × 2.5 cm geometrical shapes (square or diamond) mapped to either a left or a right button press (counterbalanced across participants). Participants were instructed to press the button associated with the stimulus, regardless of where it appeared. There were 380 trials, 190 congruent (side of presentation and required response the same) and 190 incongruent (required response opposite to the side of presentation), presented in random order. **(A)** Grand average over the electrode cluster surrounding Cz and **(B)** over a cluster chosen following a data-driven approach.

#### Cognitive assessment

Cognitive ability was assessed with the Brief Cognitive Ability Measure (B-CAM), a computerized battery developed to assess domains typically affected in people with HIV, including processing speed, attention and cognitive control (Flanker task, Corsi block task), episodic memory (verbal list learning), and phonemic fluency (FAS). Performance on these tests reflects a single underlying construct in people with HIV, so they can be combined to produce a single, continuous measure of cognitive ability (M. J. Brouillette et al., 2019; Koski et al., 2011). Lower B-CAM scores have been shown to relate to cognitive symptoms, and to a reduced ability to carry out work and social roles (Mayo et al., 2019a). 53 participants included here also took part in a separate sub-study that administered standard neuropsychological testing, covering six domains, at least two tests per domain (Antinori et al., 2007).

#### Co-morbidity characterization

Self-reported depression and anxiety symptoms were assessed with the Hospital Anxiety and Depression Scale (Zigmond & Snaith, 1983). Presence of metabolic syndrome was calculated as a binary variable (present/absent)in accordance with the National Cholesterol Education Program Adult Treatment Panel (ATP) III guidelines (Lipsy, 2003).

#### Electroencephalographic Recordings and Analysis

EEG was recorded with a 256-channel HydroCel Geodesic-Sensor-Net (Electrical Geodesics, Inc., Eugene, OR) using Net Station 5 software. Data were collected using the electrode Cz as reference and keeping electrode impedance below 50 kΩ. Seventy-eight electrodes located over the neck and cheeks were excluded before data pre-processing, as these were consistently contaminated with muscle artifacts.

EEG data were analyzed using Brainstorm (Tadel, Baillet, Mosher, Pantazis, & Leahy, 2011). Pre-processing followed standard procedures as recommended by Luck (2014)(Luck, 2014) and Brainstorm. Continuous EEG recordings were filtered (0.1-30 Hz) down-sampled to 500 Hz and re-referenced to the right and left mastoid electrodes. Automatic blink detection was done using four electrodes located above and below each eye and artifact correction was performed with Signal-Space Projection.

Only correct trials without activity exceeding ± 100 μV were analyzed. The average number of trials included for the Simon task was 109 (SD 34) (congruent), and 102 (SD 38) (incongruent). For the oddball task, a mean of 51 (SD 8) large deviant and 49 (SD 9) small deviant trials were included.

#### ERP analysis

Time windows were defined according to the maxima and time distributions of grand average waveform of the conditions of interest for each task. ERPs of interest for the oddball task were the N100 (mean amplitude between 90 and 120ms) and the P300 (mean amplitude between 280 - 400ms). For the Simon task, the analysis focused on the N200 (mean amplitude between 200-300 ms) and the P300 (mean amplitude between 300 - 550 ms). All epochs had a 200 ms pre-stimulus baseline.

Data extraction was twofold. First, clusters of electrodes centered on electrodes at which ERPs have a maximum distribution were used. The amplitude of the P300 was extracted at a cluster of electrodes (E081, E045, E132) centered on Cz (Melara, Wang, Vu, & Proctor, 2008; John Polich, 2012). The amplitude of the N100 evoked by the oddball task was extracted from the cluster centered at Cz, and the N200 evoked by the Simon task at a cluster centered on frontocentral electrodes (E015 and E023) (Folstein & Van Petten, 2008). Additional extraction was in line with previous studies showing decreased P300 amplitude in cognitively impaired participants (Gu et al., 2018; Tsolaki et al., 2017). This was focused on clusters located on parieto-occipital and central-parietal at clusters (**Figures 2** and **3**) with maximum differences between conditions (corrected with False Discovery Rate at the 0.01 level).

#### MRI acquisition and analysis

MRI was acquired using a 3T Siemens Tim TRIO v17 scanner (Siemens AG, Erlangen, Germany) with a 12-channel transmit/receive head coil. The scanning protocol included a T1-weighted three-dimensional magnetization-prepared rapid acquisition gradient echo sequence (repetition time (TR)/echo time (TE)/inversion time (TI)=2300/2.98/900ms; voxel=1.0mm^3^; flip angle=9°). The T1-weighted data was processed, denoised and registered as described in Sanford et al., (2017). After registration, brain regions identified on the ICBM152 template (Fonov, Evans, McKinstry, Almli, & Collins, 2009) were mapped back to the subject’s data to accurately identify specific structures of interest. The volumes in each of these regions were defined as the volume (cm^3^) of all segmented voxels in the standard space.

#### Head Modeling and ERP Source Estimation

Source estimation was carried out for participants who underwent MRI. FreeSurfer (http://surfer.nmr.mgh.harvard.edu/) was used to extract and register cortical surfaces and white matter envelopes. Head modeling was performed using a symmetric boundary element method as implemented in the OpenMEEG software (Gramfort, Papadopoulo, Olivi, & Clerc, 2010). Minimum-norm imaging was used to estimate cortical sources of ERPs (Baillet, Mosher, & Leahy, 2001). Current density maps and dipole orientations constrained to be normal to cortex were used for inverse modeling. Z-score transformations were done to convert current density values at the subject level to a score representing the number of standard deviations with respect to the baseline period. Sources were rectified to absolute values and projected to default anatomy template before averaging across participants. Gaussian smoothing with a 3 mm full width at half maximum was applied to the final averages.

#### Statistical Analyses

Statistical analyses were performed using R version 3.4.2 (R Core Team, 2018). Repeated measures ANOVA with trial type as within subject factor was used to test for the effects of trial type on the task performance and ERPs of interest (N100, N200 and P300). This was done independently for each task using the factors small vs. large deviant vs. standard for the oddball task and congruent vs. incongruent for the Simon task. For the P300 effects - evaluated at two electrodes-the additional factor electrode was included. To address our main question on the neural mechanisms underlying cognitive ability variation in HIV+ individuals, we performed multiple linear regressions assessing the contribution of ERPs amplitude in predicting overall cognitive ability (B-CAM).

Similarly, the effects of current and nadir CD4 cell counts on electrophysiological variables were assessed using multiple linear regression. Given that most participants had current CD4 cell counts in the normal range, and in line with recent MRI literature in cART-treated patients, it was hypothesized that the nadir CD4 count would be more relevant in predicting HIV-related EEG effects. Age and education (dichotomized as less than university, or at least some university education) were included in all models.

Pearson correlations were used to test whether volumes of subcortical nuclei previously shown to be affected in HIV and known to be involved in the cognitive tasks studied here explained the variance observed in P300 amplitudes (thalamus, globus pallidus and putamen). Correlations between P300 amplitude and extra-cerebral cerebrospinal fluid (CSF) and total grey matter volumes were examined to determine if diminished amplitudes were explained by overall brain volume or non-specific differences in conductance due to expanded CSF space between the cortex and scalp.

## Results

### Participant characteristics

Demographic and clinical variables are summarized in **Table 1**. All participants were taking cART and had intact hearing in the frequency range of the stimuli used in the oddball task.

**Table 1:** Demographic, clinical, and neuropsychological characteristics of the participants, (N=66). BCAM: Brief Cognitive Ability Measure; ANI: Asymptomatic Neuropsychological Impairment; MND: mild neurocognitive disorder; HADS Hospital Anxiety and Depression Scale; IQR: inter-quartile range; SD: standard deviation.

| Table 1: Demographic, clinical, and neuropsychological characteristics of the participants (N=66). |  |
| --- | --- |
| Characteristic |  |
| Age, mean (SD), y | 54 (7) |
| Education, No. (%) |  |
| College diploma or less | 39 (59%) |
| Any university education | 27 (41%) |
| Cognitive Assessment |  |
| B-CAM score, mean (SD) | 20.7 (4.2) |
| Neuropsychological HAND diagnosis |  |
| Impaired (ANI or MND), No. (%) | 30 (56) |
| Anxiety (HADS) | 7.0 (4.5) |
| Depression (HADS) | 4.8 (4.0) |
| Presence of metabolic syndrome, No. (%) | 29 (44%) |
| Current CD4 cell count, median (IQR), cells/μL | 607 (477 - 852) |
| Nadir CD4 cell count, median (IQR), cells/μL | 170 (109-241) |
| Plasma Viral Load |  |
| Virologically suppressed (<50 copies/mL), No. (%) | 63 (95%) |
| Duration of HIV infection, mean (SD), y | 18 (7) |

### Simon and oddball task performance

In the oddball task, large deviants were discriminated more accurately and faster than small deviants. Repeated measures ANOVA with trial type as within-subject factor showed a significant effect of trial type on RT (F (2, 52) = 60.79, p < 0.001, ε = 0.70, and d’ (F (1, 53) = 14.06, p < 0.001, ε = 0.21). Post hoc Bonferroni-corrected pairwise comparisons showed that RTs were faster for large vs. small deviant (t (53) = −7.52, p < 0.001), and vs. standard trials (t (53) = −10.78, p < 0.001), as well as for standard vs. small deviants (t (53) = 4.51, p < 0.001). For the Simon Task, trial type had significant effects on reaction time (F (1, 57) = 80.56, p < 0.001, ε = 0.58) and accuracy (F (1, 57) = 39.99, p < 0.00, ε = 0.41), with participants less accurate and slower on incongruent trials.

### ERPs in response to Simon and oddball tasks

In the oddball task, the factors trial type (F (2, 53) = 42.51, p < 0.0001, ε = 0.45), cluster of electrodes (F (1, 53) = 20.48, p < 0.0001, ε = 0.28), and their interaction (F (2, 53) = 11.11, p < 0.0001, ε = 0.17), had significant effects on the amplitude of the P300. Post-hoc pairwise comparisons (Bonferroni-corrected) showed that the amplitude of the P300 for the large deviant was significantly larger from that of the small deviant and the standard tone (*p* < 0.0001). The effect of trial type was also significant on the N100 (F (2, 53) = 18.87, p < 0.0001, ε = 0.26), with the N100 evoked by the large deviant significantly different from the small deviant and standard tone (*p*< 0.0001). Evoked responses for the oddball task are shown in **Figure 2.**

In the Simon task, neither trial type nor cluster of electrodes had significant effects on the P300 or N200 amplitude. A significant interaction was found between these two factors (F (1, 58) = 9.40, p = 0.003, ε = 0.14), but post-hoc pairwise comparisons showed no significant differences between the P300 amplitude of congruent and incongruent trials. Evoked responses for the Simon task are shown in **Figure 3.**

### Relationship between task performance and cognitive ability

The validity of both tasks as indicators of overall cognitive ability was confirmed using multiple linear regression to predict overall cognitive ability (B-CAM scores). In the oddball task, *d’* for the large deviant and age explained variation in B-CAM scores (F (3,50) = 10.3, p < 0.0001). Neither RTs nor *d’* for the small deviant predicted cognitive ability. Thus, only large deviant trials were considered for further analysis.

In the Simon task, a significant relationship was found between incongruent trial RT and cognitive ability (F (3,55) = 4.56, p = 0.006). Neither congruent RT nor accuracy predicted B-CAM scores. Hence, the incongruent condition was retained for subsequent analysis.

### Relationships between task performance and HIV severity

Multiple linear regression showed that neither CD4 cell count (nadir, current), age nor education predicted performance of either task (all *p* > 0.1).

### Relationships between ERPs and cognitive ability

Multiple linear regression models were fitted for each task to evaluate if neural activity reflecting specific cognitive processes related to overall cognitive ability. B-CAM score was not significantly predicted by the models when the amplitudes of early ERPs (N100 and N200) were included (*p* > 0.1). In contrast, later ERPs were associated with overall cognitive ability in both tasks. The amplitude of the P300 evoked by the oddball task at the cluster of electrodes around Pz was significantly related to B-CAM (F (3, 50) = 4.73, p = 0.005), with a marginal effect of P300 amplitude (*p* = 0.1) and age (*p* = 0.04). At the central parietal cluster, the relationship was also significantly related to B-CAM (F (3, 50) = 6.60, p = 0.0007), (**Table 2**). In the Simon task, the relationship was similar at the cluster around Cz (F (3, 55) = 5.32, p = 0.002) and replicated at the second parieto-occipital cluster of electrodes: B-CAM score was predicted by the amplitude of the P300 (p=0.0007) and age (p= 0.008) (F (3, 55) = 8.02, p = 0.0001). (**Table 2**).

**Table 2:**
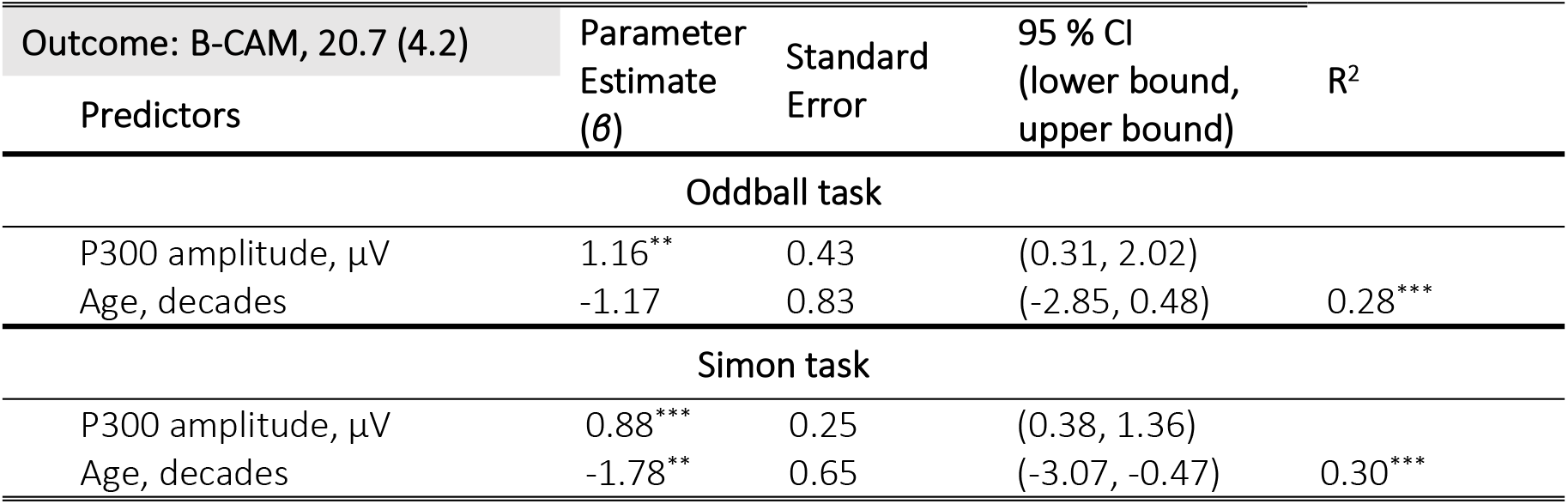
Multiple linear regressions predicting cognitive ability (BCAM score). To facilitate interpretation age is expressed in decades. All regressions were corrected for education. \**p* < 0.05; ** *p* < 0.01; *** *p* < 0.001

### Relationships between ERPs and immunocompromise

For each of the tasks, linear regressions were carried out to test if current or past HIV infection severity contributed to the variation in neural activity in early or late stages of cognitive processing. The amplitudes of early ERPs were not significantly explained by nadir or current CD4 count. Nadir CD4 counts (*p* = 0.0006) and age (*p* = 0.0001) independently contributed to predicting the P300 amplitude evoked by the oddball task at the temporal cluster of electrodes, with an increase in 100 cells/μL in the nadir CD4 count related to an increase in amplitude of about 1 μV, (F (3,50) = 11.81, p < 0.000001). A similar significant contribution was found at the cluster centered on Pz (*p*_*age*_ = 0.007, *p*_*nadir_cd4*_ = 0.05). Adding current CD4 count accounted only for an additional 3% of the variance observed in the P300 amplitude at the temporal cluster of electrodes. Neither nadir nor current CD4 counts explained the amplitude of the P300 evoked by the Simon task at either cluster of interest (*p* > 0.9) (**Table 3**).

**Table 3:**
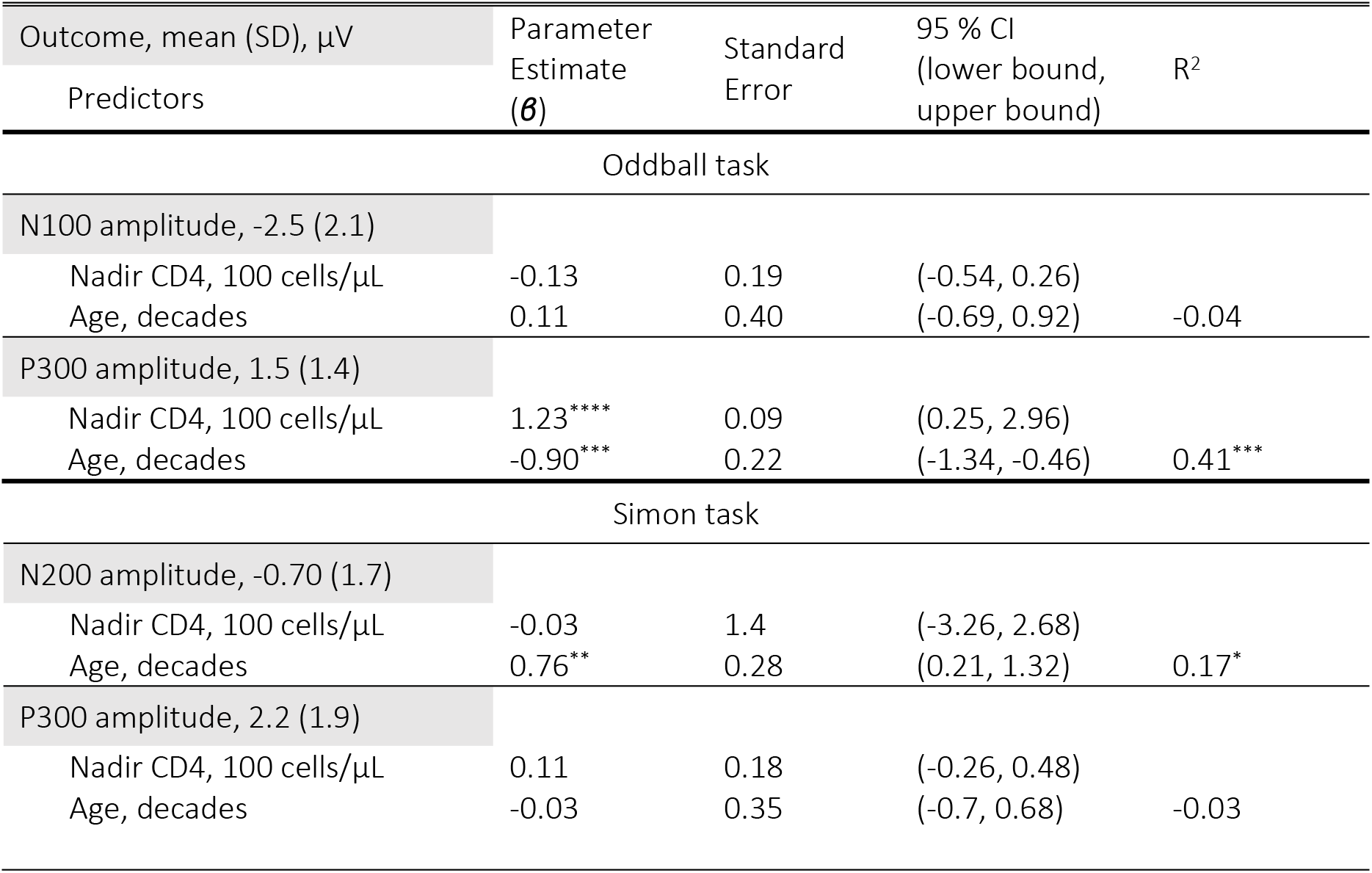
Multiple linear regressions predicting ERP amplitudes. To facilitate interpretation, age is expressed in decades, and nadir CD4 count in 100 cells/μL. All regressions were corrected for education (college or less, or any university). * *p* < 0.05; ** *p* < 0.01; *** *p* < 0.0001.

We repeated the linear regressions restricted to the participants who had completed both tasks (n = 47) to ensure that the observed differences in the relationship between nadir CD4 count and P300 amplitudes in the two tasks was not due to differences in the participants included in each analysis. The pattern of results remained unchanged; *p* >0.9 for the Simon task and *p* = 0.0001 for the oddball task. We additionally tested if depression or anxiety as measured with the HADS, or presence of metabolic syndrome explained any variance in the ERPs of interest. None of the regressions was statistically significant.

### ERP source localization

**Figure 4** shows the source estimations for both tasks.

**Figure 4:**
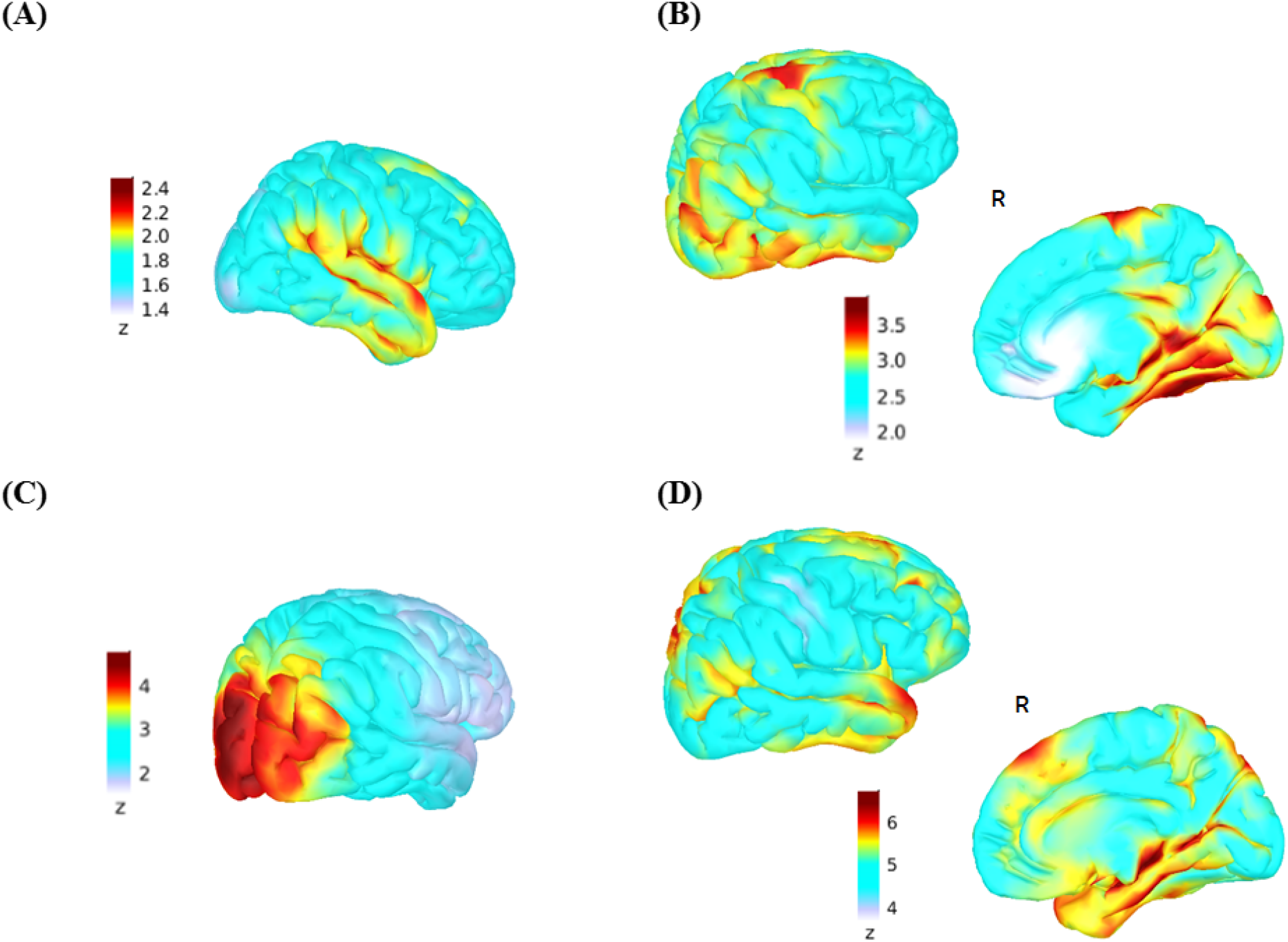
Source Estimation for EEG activity in response to the oddball (A, B) and Simon tasks (C, D). Current density maps for both tasks were normalized in relation to values during the baseline period prior to stimulus presentation (−200 to 0 ms). Z-scores are with respect to the baseline period. **(A)** Current density maps during early auditory processing in response to the large deviant tone between 70 and 90 ms post stimulation. **(B)** Current density maps indicating sources for the P300 peak activations (280 - 400 ms) in response to the large deviant tone. The P300 evoked during the oddball task showed current density foci peaking in middle and inferior temporal gyri, posterior cingulate, temporal-parietal junction and the posterior superior frontal gyrus, principally on the right. **(C)** Early visual activity between 90 - 120 ms in response to incongruent stimuli. **(D)** Peak activations during the P300 time window in response to incongruent stimuli (300 - 550 ms). Lateralization to the right hemisphere also occurred for the P300 of the Simon task. As in the oddball task, activity localized to the temporal-parietal junction, but otherwise engaged different cortical regions: the anterior cingulate, middle and anterior-superior frontal gyrus and superior temporal gyrus. R= right hemisphere.

### Subcortical contributions to P300 amplitude

Thalamus volumes were related to the amplitude of the P300 evoked by the oddball task (oddball P300, *n* =27, *r* = 0.36 *p* = 0.03) but not with the P300 evoked by the Simon task (*p’s* > 0.14). The volume of the globus pallidus showed similar relationships with the P300 amplitude of both tasks (oddball P300, *n* = 27: *r* = 0.32, *p* = 0.03, Simon P300, *r* = 0.46, *p* = 0.008). These were not general effects: there was no correlation between P300 amplitude evoked by either task and putamen volumes (**Fig. 5**), nor was it related to total grey matter volume, or extra-cerebral CSF volume (*p*s > 0.2).

**Figure 5:**
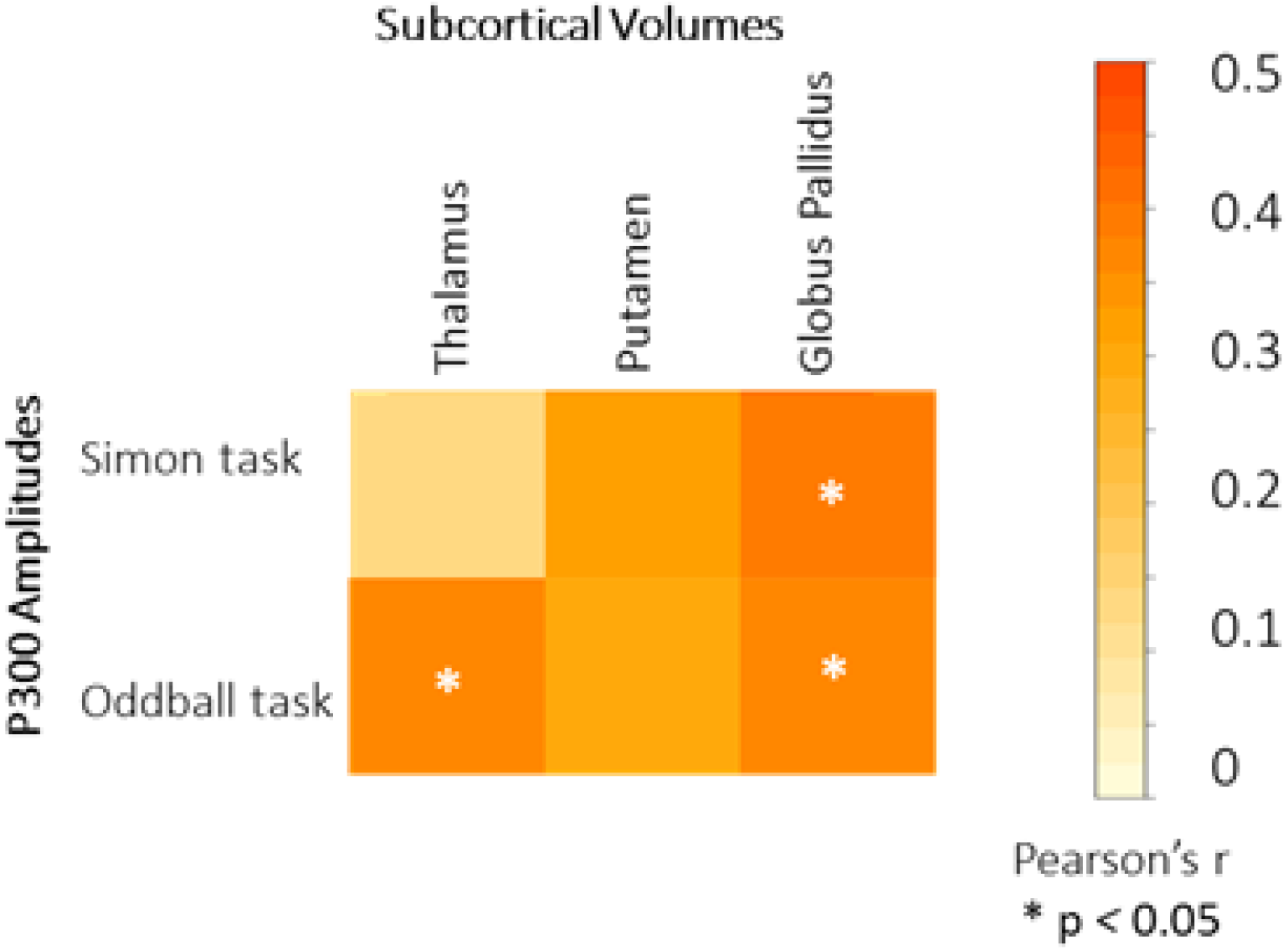
Relationships between subcortical volumes and P300 amplitudes in the Simon and oddball tasks. Asterisks indicate significant correlations (uncorrected). The scale indicates Pearson’s r coefficients.

## Discussion

This study of the neural mechanisms underlying variation in cognitive ability in older men with well-controlled HIV found evidence for specific vulnerabilities in processing stages and neural circuits. Behavioral measures from two tasks differing in higher-order response control and sensory-attentional requirements were related to overall cognitive ability in this large, well-characterized, non-demented sample. Stimulus-locked EEG activity evoked by these tasks provided evidence that early sensory and cognitive processing stages in both tasks did not relate to overall cognitive ability, nor was this activity affected by past HIV severity indicated by nadir CD4 count. In contrast, the amplitudes of the P300 waveforms evoked in both tasks did relate to cognitive ability. In the auditory oddball task, amplitudes also were related to nadir CD4 count. Co-morbid depression, anxiety and metabolic syndrome did not explain variation in P300 amplitude. These ERP effects were not due to global brain atrophy. Instead, we found preliminary MRI evidence implicating subcortical nuclei in P300 amplitude variation.

Prior small studies have compared early potentials evoked by auditory oddball and visual stimulation tasks in HIV+ and HIV– samples. Most reported no difference in N100 or N200 amplitudes between groups (Chao, Lindgren, Flenniken, & Weiner, 2004; J Polich, 2000; Tartar et al., 2004), although two studies found reduced N200 amplitudes in HIV+ groups (J Polich, 2000; Tartar, McIntosh, Rosselli, Widmayer, & Nash, 2014). Here, in a larger sample, using cortical source estimation we show that these earlier processing stages in auditory and visual modalities are relatively spared. Some evidence from structural and functional MRI studies also supports this claim (Chang & Shukla, 2018; Masters & Ances, 2014).

Cognitive ability was related to the amplitude of the P300 evoked by both tasks. Past work on the P300 in HIV, mainly evoked with oddball tasks, reported amplitude reductions for HIV+ compared to HIV– controls in smaller samples with shorter duration of HIV infection than those we studied (reviewed in Fernández-Cruz & Fellows, 2017). Here, we confirm the relevance of the P300 as an indicator of brain dysfunction in HIV, showing for the first time that it relates to cognitive ability in this population, and finding that this relationship is present across two different tasks in the same sample. While smaller ERP amplitudes are a well-known feature of aging (Gajewski, Ferdinand, Kray, & Falkenstein, 2018; Rossini, Rossi, Babiloni, & Polich, 2007), the effects reported here were over and above those related to age.

Neural activity in this time window in the oddball task reflects attention towards working memory contents (John Polich, 2007) and arises primarily from the anterior cingulate, superior temporal gyrus, temporal-parietal junction and prefrontal cortex (Walz et al., 2013). The Simon task P300 activity reflects action selection in the face of conflict, requiring inhibition of a prepotent response (Wang, Li, Zheng, Wang, & Liu, 2014), and originates principally from the middle and inferior frontal gyri and the superior parietal lobule (Xia, Li, & Wang, 2016). Our source localization findings are largely in line with this literature in healthy subjects. We found that temporal-parietal junction, inferior temporal, pre-central and posterior regions of the superior frontal gyrus as well as posterior cingulate cortex contributed to P300 activity during the oddball task. P300 sources in the Simon task included middle and superior frontal gyrus, anterior cingulate, superior temporal gyrus and temporal-parietal junction. Thus, there were shared and distinct cortical sources for activity in this time window, across tasks.

The severity of past immunosuppression, indicated by the nadir CD4 count, explained variance in P300 amplitude in the oddball task only. There are likely multiple contributors to cognitive dysfunction in HIV (Mayo et al., 2019b); this result suggests that P300 amplitude may be a good general indicator of brain health, but markers of specific neural circuit dysfunction may be better suited to detecting the specific effects of HIV infection on the brain. We provide preliminary evidence that the apparent specificity of the auditory oddball P300 may relate to thalamus dysfunction, finding that smaller thalamus volumes were associated with smaller oddball P300 amplitude. The thalamus contributes to stimulus monitoring for subsequent evaluation according to context and working memory content and is engaged in auditory and visual oddball tasks (Cacciaglia et al., 2015; Klostermann et al., 2006). Recent neuroimaging studies using MRI, fMRI and PET have shown that the thalamus is amongst the structures most affected in HIV(Sanford, Ances, et al., 2018; Sanford, Fellows, Ances, & Collins, 2018; Sanford et al., 2017; Wade et al., 2015), although this is not a universal finding (Ances, Ortega, Vaida, Heaps, & Paul, 2012; Janssen et al., 2016). Alternatively, this apparent circuit selectivity could reflect vulnerability of one or more temporal-parietal cortical regions; the limited sample with MRI data available here does not allow a strong test of this possibility. In any case, these results suggest that cognitive dysfunction in HIV extends beyond fronto-striatal regions, which have been suggested as key targets of HIV infection (Melrose, Tinaz, Castelo, Courtney, & Stern, 2008; Plessis et al., 2014). Together, these results suggest that events occurring at or triggered at the time of most-severe HIV infection (as indexed by nadir CD4 count) might continue to show an impact on the brain, even after good immune recovery (“legacy effect”). Although a prospective study would be needed to fully support this hypothesis, early treatment initiation and on-going effective viral suppression might limit or prevent brain injury in persons living with HIV. The few longitudinal studies to date show no strong evidence for progressive volume loss in cART treated individuals, at least over short periods (Cole et al., 2018; Sanford, Fellows, et al., 2018), suggesting HIV-related brain injury indeed relates to pathophysiological processes triggered before systemic viral control has been achieved.

A third explanation for this apparent circuit-specific vulnerability is that the P300 waveforms evoked by these two tasks differ in their measurement properties. While there were no ceiling or floor effects in either task, and the range and standard deviation of P300 amplitudes was similar, it could be that the signal-to-noise ratio of the Simon task-evoked P300 is lower than that of the oddball task, which would make it harder to detect a relationship with nadir CD4 count. Whether the reasons are technical or neurological, our findings argue that the auditory oddball task-evoked P300 is more suitable for detecting HIV-related brain dysfunction than the P300 evoked by the Simon task.

Attenuation of P300 amplitude in auditory oddball and other tasks has been reported in other cognitively impaired populations, such as Alzheimer's disease and amnestic mild cognitive impairment (Gu et al., 2018; Tsolaki et al., 2017), indicating that P300 amplitude reduction is not specific for HIV-associated neurocognitive disorders. EEG alone is thus better suited to detecting and quantifying brain dysfunction, rather than identifying specific underlying causes.

After accounting for age and education effects, nadir CD4 count only explains some of the variance in P300 amplitude in the oddball task, and none in the Simon task. This emphasizes the need for more comprehensive identification of the factors explaining brain dysfunction in HIV. These could include additional direct effects of HIV not indicated by nadir CD4 count, and many additional candidate mechanisms co-occurring or indirectly related to HIV status, ranging from cerebrovascular injury to mental health co-morbidities, to social factors such as stigmatization (Clifford, 2017; Lam, Mayo, Scott, Brouillette, & Fellows, 2018). Although the present study was not focused on such variables, preliminary analyses found no effect of anxiety, depression or metabolic risk (as a proxy for cerebrovascular injury) on the EEG measures in this sample.

This study has limitations. The findings can be generalized only to older, relatively well-educated men on cART without clinically obvious dementia; there is a need to carry out similar work in women with HIV. As only a subset of the sample had structural MRI, there was insufficient statistical power to carry out whole-brain analyses linking ERP and MRI; this was not an *a priori* aim of this study. Thus, the observed relationships between subcortical volumes and evoked potentials should be treated as exploratory, requiring replication in a new sample. Whole brain approaches would provide additional information on possible regional cortical or white matter contributors to the EEG variance, not tested here.

## Conclusion

These are the first EEG and source modeling results contrasting two tasks tapping different cognitive domains in the same large, well-characterized sample of men with well controlled HIV infection. We showed that evoked potentials in the P300 time window are promising markers of cognitive ability in the mildly impaired-to-normal range of neuropsychological performance, and suggest that tasks and EEG measures related to cortico-thalamic circuits may be particularly useful for probing brain health in people living with HIV. The effects of HIV at the time of untreated infection, indexed by nadir CD4 count, are detectable with EEG even after many years of cART, and specific EEG measures in turn relate to current cognitive status. There appear to be additional, unmeasured contributors to EEG variance in these older men, beyond demographic, mood and presence metabolic syndrome. It is increasingly clear that cognitive impairment in people living with HIV can have many contributors, at least some of which are likely to be treatable or remediable (Joska, Fincham, Stein, Paul, & Seedat, 2010; Lam et al., 2018; Mayo et al., 2019b).

The findings here suggest that evoked potentials, a cheap and widely available readout of neural function, may prove useful in targeting or monitoring novel treatment or rehabilitation approaches to this common and quality-of-life-limiting aspect of living with HIV.

## Authors contributions

ALFC and LKF conceived of the study and designed the experiments. AFC collected data, carried out analysis and drafted the manuscript. CMC contributed to data collection and analysis, and contributed to the manuscript. RS and DLC contributed to the MRI analysis and revised the manuscript. NEM provided advice on statistical methods. LKF, MJB and NEM conceived of the BHN study and related sub-studies, oversaw all data collection and analysis, and revised the manuscript.

## Acknowledgements

Authors would like to thank Christine Déry and Susan Scott for their help with participant recruitment and data analysis, Marcus Sefranek for technical assistance and the Positive Brain Health Now investigators, research staff and participants for making this work possible.

## References

Ances, B. M., Ortega, M., Vaida, F., Heaps, J., & Paul, R. (2012). Independent effects of HIV, aging, and HAART on brain volumetric measures. Journal of Acquired Immune Deficiency Syndromes (1999), 59(5), 469–477.

Antinori, A., Arendt, G., Becker, J. T., Brew, B. J., Byrd, D. A., Cherner, M., ... Wojna, V. E. (2007). Updated research nosology for HIV-associated neurocognitive disorders. Neurology, 69, 1789–1799.

Baillet, S., Mosher, J. C., & Leahy, R. M. (2001). Electromagnetic brain mapping. IEEE Signal Processing Magazine, 18(6), 14–30.

Becker, B. W., Thames, A. D., Woo, E., Castellon, S. A., & Hinkin, C. H. (2011). Longitudinal change in cognitive function and medication adherence in HIV-infected adults. AIDS and Behavior, 15(8), 1888–1894.

Brouillette, M.-J., Mayo, N. E., Fellows, L. K., Koski, L., & Cysique, L. A. (n.d.). Sensitivity and specificity of a brief cognitive test battery for mild HAND. In Preparation.

Brouillette, M. J., Fellows, L. K., Finch, L., Thomas, R., & Mayo, N. E. (2019). Properties of a brief assessment tool for longitudinal measurement of cognition in people living with HIV. PLoS ONE, 14(3), 1–13.

Cacciaglia, R., Escera, C., Slabu, L., Grimm, S., Sanjuán, A., Ventura-Campos, N., & Ávila, C. (2015). Involvement of the human midbrain and thalamus in auditory deviance detection. Neuropsychologia, 68, 51–58.

Chang, L., & Shukla, D. K. (2018). Imaging studies of the HIV-infected brain. Handbook of clinical neurology (1st ed., Vol. 152). Elsevier B.V.

Chao, L. L., Lindgren, J. a, Flenniken, D. L., & Weiner, M. W. (2004). ERP evidence of impaired central nervous system function in virally suppressed HIV patients on antiretroviral therapy. Clin Neurophysiol, 115(7), 1583–1591.

Clifford, D. B. (2017). HIV-associated neurocognitive disorder. Current Opinion in Infectious Diseases, 30(1), 117–122.

Cole, J. H., Caan, M. W. A., Underwood, J., De Francesco, D., van Zoest, R. A., Wit, F. W. N. M., ... Sharp, D. J. (2018). No Evidence for Accelerated Aging-Related Brain Pathology in Treated Human Immunodeficiency Virus: Longitudinal Neuroimaging Results From the Comorbidity in Relation to AIDS (COBRA) Project. Clinical Infectious Diseases, 66(June), 1899–1909.

Fernández-Cruz, A. L., & Fellows, L. K. (2017). The electrophysiology of neuroHIV: A systematic review of EEG and MEG studies in people with HIV infection since the advent of highly-active antiretroviral therapy. Clinical Neurophysiology, 128(6), 965–976.

Folstein, J. R., & Van Petten, C. (2008). Influence of cognitive control and mismatch on the N2 component of the ERP: A review. Psychophysiology, 45(1), 152–170.

Fonov, V., Evans, A., McKinstry, R., Almli, C., & Collins, D. (2009). Unbiased nonlinear average age-appropriate brain templates from birth to adulthood. NeuroImage, 47, S102.

Gajewski, P. D., Ferdinand, N. K., Kray, J., & Falkenstein, M. (2018). Understanding Sources of Adult Age Differences in Task Switching: Evidence from Behavioral and ERP Studies. Neuroscience & Biobehavioral Reviews, 92(June), 255–275.

Goodkin, K., Miller, E. N., Cox, C., Reynolds, S., Becker, J. T., Martin, E., ... Sacktor, N. C. (2017). Effect of ageing on neurocognitive function by stage of HIV infection: evidence from the Multicenter AIDS Cohort Study. The Lancet HIV, 4(9), 411–422.

Gramfort, A., Papadopoulo, T., Olivi, E., & Clerc, M. (2010). OpenMEEG: Opensource software for quasistatic bioelectromagnetics. BioMedical Engineering Online, 9, 1–20.

Groff, B. R., Wiesman, A. I., Rezich, M. T., O’Neill, J., Robertson, K. R., Fox, H. S., ... Wilson, T. W. (2020). Age-related visual dynamics in HIV-infected adults with cognitive impairment. Neurology - Neuroimmunology Neuroinflammation, 7(3), e690.

Gu, L., Chen, J., Gao, L., Shu, H., Wang, Z., Liu, D., ... Zhang, Z. (2018). Cognitive reserve modulates attention processes in healthy elderly and amnestic mild cognitive impairment: An event-related potential study. Clinical Neurophysiology, 129(1), 198–207.

Heaton, R. K., Clifford, D. B., Franklin, D. R., Woods, S. P., Ake, C., Vaida, F., ... Grant, I. (2010). HIV-associated neurocognitive disorders persist in the era of potent antiretroviral therapy: CHARTER Study. Neurology, 75, 2087–2096.

Heaton, R. K., Marcotte, T. D., Mindt, M. R., Sadek, J., Moore, D. J., Bentley, H., ... Grant, I. (2004). The impact of HIV-associated neuropsychological impairment on everyday functioning. Journal of the International Neuropsychological Society : JINS, 10(3), 317–331.

Holt, J. L., Kraft-Terry, S. D., & Chang, L. (2012). Neuroimaging studies of the aging HIV-1-infected brain. Journal of Neurovirology, 18, 291–302.

Janssen, M. A. M., Hinne, M., Janssen, R. J., van Gerven, M. A., Steens, S. C., Góraj, B., ... Kessels, R. P. C. (2016). Resting-state subcortical functional connectivity in HIV-infected patients on long-term cART. Brain Imaging and Behavior.

Joska, J. A., Fincham, D. S., Stein, D. J., Paul, R. H., & Seedat, S. (2010). Clinical correlates of HIV-associated neurocognitive disorders in South Africa. AIDS and Behavior, 14(2), 371–378.

Klostermann, F., Wahl, M., Marzinzik, F., Schneider, G. H., Kupsch, A., & Curio, G. (2006). Mental chronometry of target detection: Human thalamus leads cortex. Brain, 129(4), 923–931.

Koski, L., Brouillette, M. J., Lalonde, R., Hello, B., Wong, E., Tsuchida, A., & Fellows, L. K. (2011). Computerized testing augments pencil-and-paper tasks in measuring HIV-associated mild cognitive impairment. HIV Medicine, 12(8), 472–480.

Lam, A., Mayo, N. E., Scott, S., Brouillette, M.-J., & Fellows, L. K. (2018). HIV-Related Stigma Affects Cognition in Older Men Living with HIV. JAIDS Journal of Acquired Immune Deficiency Syndromes, 1.

Lew, B. J., McDermott, T. J., Wiesman, A. I., O’Neill, J., Mills, M. S., Robertson, K. R., ... Wilson, T. W. (2018). Neural dynamics of selective attention deficits in HIV-associated neurocognitive disorder. Neurology, 91(20), e1860–e1869.

Lipsy, R. J. (2003). The National Cholesterol Education Program Adult Treatment Panel III Guidelines. Journal of Managed Care Pharmacy, 9(1 Supp A), 2–5.

Luck, S. J. (2014). Basic principles of ERP recording. An Lntroduction to the Event-Related Potential Technique, 147–183.

Luck, S. J., Woodman, G. F., & Vogel, E. K. (2000). Event-related potential studies of attention. Trends in Cognitive Sciences, 4(11), 432–440.

Masters, M. C., & Ances, B. M. (2014). Role of neuroimaging in HIV-associated neurocognitive disorders. Semin.Neurol., 34(1098-9021 (Electronic)), 89–102.

Mayo, N. E., Brouillette, M.-J., & Fellows, L. K. (2016). Understanding and optimizing brain health in HIV now: Protocol for a longitudinal cohort study with multiple randomized controlled trials. BMC Neurology, 16:8.

Mayo, N. E., Brouillette, M.-J., Scott, S. C., Harris, M., Smaill, F., Smith, G., ... Fellows, L. K. (2019a). Relationships between cognition, function, and quality of life among HIV+ Canadian men. Quality of Life Research.

Mayo, N. E., Brouillette, M.-J., Scott, S. C., Harris, M., Smaill, F., Smith, G., ... Fellows, L. K. (2019b). Relationships between cognition, function, and quality of life among HIV+ Canadian men. Quality of Life Research, (0123456789).

Melara, R. D., Wang, H., Vu, K. P. L., & Proctor, R. W. (2008). Attentional origins of the Simon effect: Behavioral and electrophysiological evidence. Brain Research, 1215, 147–159.

Melrose, R. J., Tinaz, S., Castelo, J. M. B., Courtney, M. G., & Stern, C. E. (2008). Compromised fronto-striatal functioning in HIV: an fMRI investigation of semantic event sequencing. Behavioural Brain Research, 188(2), 337–347.

Nolan, R., & Gaskill, P. J. (2019). The role of catecholamines in HIV neuropathogenesis. Brain Research, 1702, 54–73.

Du Plessis, S., Vink, M., Joska, J. a., Koutsilieri, E., Stein, D. J., & Emsley, R. (2014). HIV infection and the fronto–striatal system. Aids, 28, 803–811.

Polich, J. (2000). Neuroelectric assessment of HIV: EEG, ERP, and viral load. International Journal of Psychophysiology, 38(1), 97–108.

Polich, John. (2007). Updating P300: An integrative theory of P3a and P3b. Clin Neurophysiol, 118, 2128–2148.

Polich, John. (2012). Neuropsychology of P300. (E. S. Kappenman & L. Steven J., Eds.), Oxford Handbook of Event-Related Potential Components. Oxford University Press.

Price, R., & Brew, B. (1988). The AIDS dementia complextle. J Infect Dis., 158(5), 1079–1083.

R Core Team. (2018). R: A Language and Environment for Statistical Computing. R Foundation for Statistical Computing, Vienna, Au(https://www.R-project.org).

Rossini, P. M., Rossi, S., Babiloni, C., & Polich, J. (2007). Clinical neurophysiology of aging brain: From normal aging to neurodegeneration. Progress in Neurobiology, 83, 375–400.

Sanford, R., Ances, B. M., Meyerhoff, D. J., Price, R. W., Fuchs, D., Zetterberg, H., ... Collins, D. L. (2018). Longitudinal Trajectories of Brain Volume and Cortical Thickness in Treated and Untreated Primary HIV Infection. Clinical Infectious Diseases, l(May), 1–23.

Sanford, R., Fellows, L. K., Ances, B. M., & Collins, D. L. (2018). Association of Brain Structure Changes and Cognitive Function With Combination Antiretroviral Therapy in HIV-Positive Individuals. JAMA Neurology, 75(1), 72.

Sanford, R., Fernandez Cruz, A. L., Scott, S. C., Mayo, N. E., Fellows, L. K., Ances, B. M., & Collins, D. L. (2017). Regionally Specific Brain Volumetric and Cortical Thickness Changes in HIV-infected Patients in the HAART era. J Acquir Immune Defic Syndr, 74, 563–570.

Sanford, R., Fernandez Cruz, A. L., Tanenbaum, A., Westerhaus, E., Nelson, B., Fellows, L. K., ... Collins, D. L. (2016). Regionally Specific Brain Volumetric and Cortical Thickness Changes in HIV+ Patients in the HAART era, Submited,.

Su, T., Schouten, J., Geurtsen, G. J., Wit, F. W., Stolte, I. G., Prins, M., ... Mulder, W. M. C. (2015). Multivariate normative comparison, a novel method for more reliably detecting cognitive impairment in HIV infection. Aids, 29(5), 547–557.

Tadel, F., Baillet, S., Mosher, J. C., Pantazis, D., & Leahy, R. M. (2011). Brainstorm: A user-friendly application for MEG/EEG analysis. Computational Intelligence and Neuroscience, 2011.

Tartar, J. L., McIntosh, R. C., Rosselli, M., Widmayer, S. M., & Nash, A. J. (2014). HIV-positive females show blunted neurophysiological responses in an emotion-attention dual task paradigm. Clin Neurophysiol, 125, 1164–1173.

Tartar, J. L., Sheehan, C. M., Nash, A. J., Starratt, C., Puga, A., & Widmayer, S. (2004). ERPs differ from neurometric tests in assessing HIV-associated cognitive deficit. NeuroReport, 15(10), 1675–1678.

Thompson, P. M., & Jahanshad, N. (2015). Novel Neuroimaging Methods to Understand How HIV Affects the Brain. Current HIV/AIDS Reports, 12, 289–298.

Tsolaki, A. C., Kosmidou, V., Kompatsiaris, I. (Yiannis), Papadaniil, C., Hadjileontiadis, L., Adam, A., & Tsolaki, M. (2017). Brain source localization of MMN and P300 ERPs in mild cognitive impairment and Alzheimer’s disease: a high-density EEG approach. Neurobiology of Aging, 55, 190–201.

Valcour, V., Shikuma, C., Shiramizu, B., Watters, M., Poff, P., Selnes, O., ... Sacktor, N. (2004). Higher frequency of dementia in older HIV-1 individuals: The Hawaii aging with HIV-1 cohort. Neurology, 63(5), 822–827.

Wade, B. S. C., Valcour, V. G., Wendelken-Riegelhaupt, L., Esmaeili-Firidouni, P., Joshi, S. H., Gutman, B. A., & Thompson, P. M. (2015). Mapping abnormal subcortical brain morphometry in an elderly HIV + cohort. NeuroImage: Clinical, 9, 564–573.

Walz, J. M., Goldman, R. I., Carapezza, M., Muraskin, J., Brown, T. R., & Sajda, P. (2013). Simultaneous EEG-fMRI Reveals Temporal Evolution of Coupling between Supramodal Cortical Attention Networks and the Brainstem. Journal of Neuroscience, 33(49), 19212–19222.

Wang, K., Li, Q., Zheng, Y., Wang, H., & Liu, X. (2014). Temporal and spectral profiles of stimulus-stimulus and stimulus-response conflict processing. NeuroImage, 89, 280–288.

Wiesman, A. I., O’Neill, J., Mills, M. S., Robertson, K. R., Fox, H. S., Swindells, S., & Wilson, T. W. (2018). Aberrant occipital dynamics differentiate HIV-infected patients with and without cognitive impairment. Brain, 141(6), 1678–1690.

Xia, T., Li, H., & Wang, L. (2016). Implicitly strengthened task-irrelevant stimulus-response associations modulate cognitive control: Evidence from an fMRI study. Human Brain Mapping, 37(2), 756–772.

Zigmond, A. S., & Snaith, R. P. (1983). The Hospital Anxiety and Depression Scale. Acta Psychiatrica Scandinavica, 67(6), 361–370.

